# Identification of Chlorophyll a-b Binding Protein AB96 as a novel TGFβ1 binding agent

**DOI:** 10.1101/2020.08.20.258590

**Authors:** Steven Lynham, Fabio Grundland Freile, Natasha M Puri, Nicola O’Reilly, Graham H Mitchell, Timothy N.C. Wells, Merlin Willcox, Richard Beatson

**Author notes:** Corresponding Author Richard Beatson.

## Abstract

The discovery of compounds and proteins from plants has greatly contributed to modern medicine, especially in malaria where both quinine and artemisinin have been the cornerstone of therapeutics. Here we describe the first known plant-derived cytokine binding agent. Chlorophyll a-b binding protein AB96, present to varying levels in the leaves of all chlorophyll-containing plants, binds to active TGFβ1. Active TGFβ1 contributes to the pathology of many infectious and neoplastic diseases and therefore, by inhibiting these processes, chlorophyll a-b binding protein opens up new approaches with therapeutic potential.

## Main Text

*Vernonia amygdalina* Del. (Compositae) (VA) is used by humans and other primates for a variety of related conditions, particularly malaria, caused by plasmodium infection, and Bilharzia, caused by helminth infection. ^1,2 3^. VA is commonly used throughout sub-Saharan Africa and is normally consumed as a decoction or tea made with boiling water, although in some cases it is consumed raw. There is a large body of ethnobotanic evidence supporting this claimed activity ^2 3^. Unusually for ethnobotanic therapies, this is supported by multiple studies in preclinical models of parasite infection using standardized preparations ^4–6^. A clinical trial in human malaria patients confirmed that the extracts have significant anti-parasitic activity ^7^. The anti-infective activity has been confirmed in the purified sesquiterpene lactone, vernodalin ^2^, however this molecule has several features which do not make it a promising starting point for standard medicinal chemistry.

TGFβ1 is a highly conserved pleiotropic cytokine that is involved in multiple healthy and disease processes, including immune regulation, fibrosis, apoptosis, angiogenesis and development. TGFβ1 exists predominantly in its latent form at high concentrations *in vivo*. The latent form consists of either TGFβ1 non-covalently bound to LAP (<u>L</u>atency <u>A</u>ssociated <u>P</u>eptide) forming the small latency complex, or the small latency complex bound to LTBP1-4 (<u>L</u>atent <u>T</u>GF <u>B</u>inding <u>P</u>rotein 1-4), which forms the large latency complex ^8^. Activation of TGFβ1 can be induced through several mechanisms including acidification and heating *in vitro* or integrin engagement, thrombospondin interaction, protease cleavage, mild mechanical or pH stress *in vivo*^8^.

From an immune perspective, its latent form is largely inert. However in its active form it can a) induce regulatory T-cells (Tregs) which suppress the function of other immune cells to limit the immune response, b) induce class switching of B cells from IgG to IgA producers, with IgA inducing less inflammatory responses c) induce tolerogenic antigen presenting cells, which promote anergy / non-response to antigen and d) inhibit the polarisation of T helper (CD4) cells ^9^ resulting in a less focused immune response.

During plasmodium infection, active TGFβ1 can be seen to peak 48h after infection in sera, believed to be owing to cleavage of LAP by specific plasmodium-derived enzymes^10^. This can be seen to correlate with the induction of Tregs^11^, which facilitate plasmodium growth^12^, indeed it is worth noting at this point that individuals of Fulani heritage in West Africa frequently carry a mutation which prevents Treg induction and have reduced susceptibility to *P. falciparum* infection ^13^. In intestinal helminth infection, TGFβ1 is induced and activated by the parasite inducing a profound increase in Tregs ^14^, to such an extent that controlled infection is being trialed for a number of autoimmune diseases including Graft versus Host Disease (GvHD)^15^. Interestingly, beyond TGFβ1 induction and activation within the host, it has recently been shown that the helminth, *H. Polygyrus*, produces its own functional TGFβ1 mimetic^16^.

The immunological effects of endotoxin-free aqueous extract of VA, on human monocyte derived dendritic cells (mo-DCs) was studied. The mo-DCs were incubated with the extract for 24 hours. The supernatant from treated cells was shown to have significantly lower concentrations of TGFβ1 (Figure 1a). This suggested a specific inhibition of the TGFβ1 production pathway. However when exogenous active TGFβ1 was added, we saw the same reduction, in a similar manner to a known active TGFβ1 binder, heparin sulphate ^17^ (Figure 1b). This suggested that an agent in the extract may be binding and ‘removing’ active TGFβ1 from the culture supernatant.

**Figure 1.**
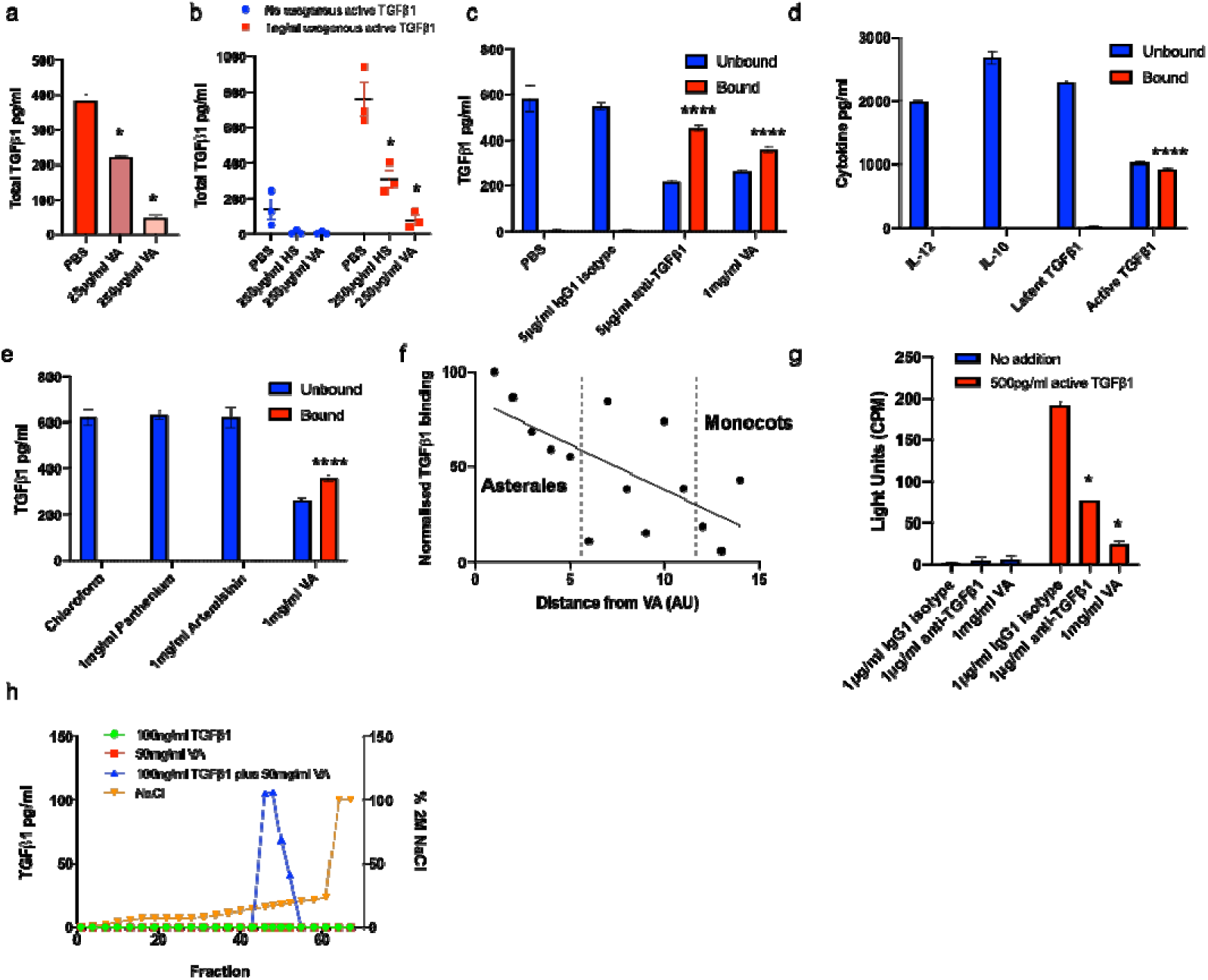
VA extract contains an active TGFβ1 neutralizing agent. The effects of VA extract on endogenous (a) and exogenous (b) TGFβ1 levels in the supernatant of monocyte derived dendritic cells. n=1 for A and n=3 for B. (c) Unbound and bound active TGFβ1 concentrations (starting: 2ng/ml) after incubation with VA extract for 2h at RT. Representative of 3 independent assays. Technical triplicates. Mean +/− SEM shown. (d) The unbound and bound IL-10, IL12, latent TGFβ1 and active TGFβ1 concentrations (starting: 4ng/ml) after incubation with VA extract for 2h at room temperature (RT). Technical triplicates. Mean +/− SEM shown. (e) The unbound and bound active TGFβ1 concentrations (starting: 2ng/ml) after incubation with sesquiterpene lactones for 2h at RT. Technical triplicates. Mean +/− SEM shown. (f) The levels of active TGFβ1 bound to aqueous extracts (1mg/ml) from different species of plants (normalized to VA extract) when incubated with 2ng/ml active TGFβ1. Species in order of x axis: 1. *Vernonia amygdalina* Del. (Compositae). 2. *Vernonia arkansana* DC. (Compositae). 3. *Helianthus annuus* L. (Compositae). 4. *Aster dumosus* Hoffm. (Compositae). 5. *Campanula lactiflora* M.Bieb. (Campanulaceae). 6. *Petroselinum crispum* (Mill.) Fuss (Apiaceae). 7. *Ocimum basilicum* L. (Lamiaceae). 8. *Solanum lycopersicum* L. (Solanaceae). 9. *Spinacia oleracea* L. (Amaranthaceae). 10. *Rosa sericea Wall.* Ex Lindl. (Rosaceae) 11. *Vitis vinifera* L. (Vitaceae). 12. *Dieffenbachia amoena* Bull. (unresolved, Araceae) 13. *Dracaena marginata* Hort. (Asparagaceae). 14. *Lilium candidum* L. (Liliaceae). Technical triplicates. Mean +/− SEM shown. (g) The effect of VA extract on luciferase activity in a functional active TGFβ1 reporter cell line. Biological triplicates. Mean +/− SEM shown. Representative of 2 independent assays. (h) The levels of TGFβ1 in different fractions eluted from an anion exchange column after loading with indicated factors, using indicated % of 2M NaCl. Unpaired student’s t-test: * p<0.05 **** p<0.0001.

To assess if VA extract was binding to active TGFβ1 we developed a simple *in vitro* method whereby we plated out the aqueous extract overnight, before incubating with active TGFβ1. The supernatant was removed for testing (the ‘unbound fraction’), before the plate was washed, acidified and neutralized, to elute the ‘bound fraction’. Active TGFβ1 was measured in the unbound and bound fractions (Figure 1c). We were concerned that this binding may be non-specific and so tested 3 other cytokines, IL-10, IL-12p70 and, importantly, latent TGFβ1, and found that this phenomenon could only be observed with active TGFβ1 (Figure 1d).

We next assessed if sesquiterpene lactones may be responsible for the observed binding, we assayed both artemisinin and parthenium finding that neither were able to bind active TGFβ1 under these conditions (Figure 1e). Subsequently, we wanted to explore if VA extract was unique in having this ability. We prepared extracts from 13 additional species, representing different branches of the plant kingdom and tested their ability to bind active TGFβ1. All species were able to bind active TGFβ1, however there was a trend suggesting that members of the Asterales order, which contains many medicinal plants, contained more of the active TGFβ1 binding compound, whilst the Monocot clade contained less. (Figure 1f). We next wished to assess if this binding rendered active TGFβ1 non-functional. Using a well-established reporter assay ^18^, we were able to show that VA extract was able to inhibit active TGFβ1 function (Figure 1g).

We next used anion exchange chromatography to show a) that binding was occurring using a different assay and b) to suggest the isoelectric point of the agent for validation purposes. As can be seen in Figure 1h, active TGFβ1 was again seen to bind to VA extract (with elution of active TGFβ1 only occurring with pre-incubation) and, in combination with the agent, active TGFβ1 eluted in fractions 46-52, which given the experimental conditions, gave the agent an isoelectric point of 4.5-5.5.

To isolate the active agent we performed a modified immunoprecipitation (IP), incubating VA extract with biotinylated active TGFβ1, or control protein and streptavidin beads. The eluant was run on a gel and whole lanes were sent for mass spectrometric analysis. Two peptides were obtained, EVIHSRWAMLGALGCVFPELLSR and FGEAVWFK which are both present in the same protein; chlorophyll a-b binding protein (Ca-bBP) AB96 (P04159). The initial hits were identified as being part of the sequence from *P. sativum* L. (Leguminosae), but the protein is well conserved and, with the sequence for VA unavailable at this current time, it is assumed that this sequence is also present in VA. These peptides were seen to have bound active TGFβ1 and not the controls (Figure 2a-c). Using *E.coli* derived recombinant full-length folded Ca-bBP AB96 from *P. sativum* L. (Leguminosae), we repeated the active TGFβ1 binding assays finding that Ca-bBP AB96 was able to bind human active TGFβ1 (Figure 2d).

**Figure 2.**
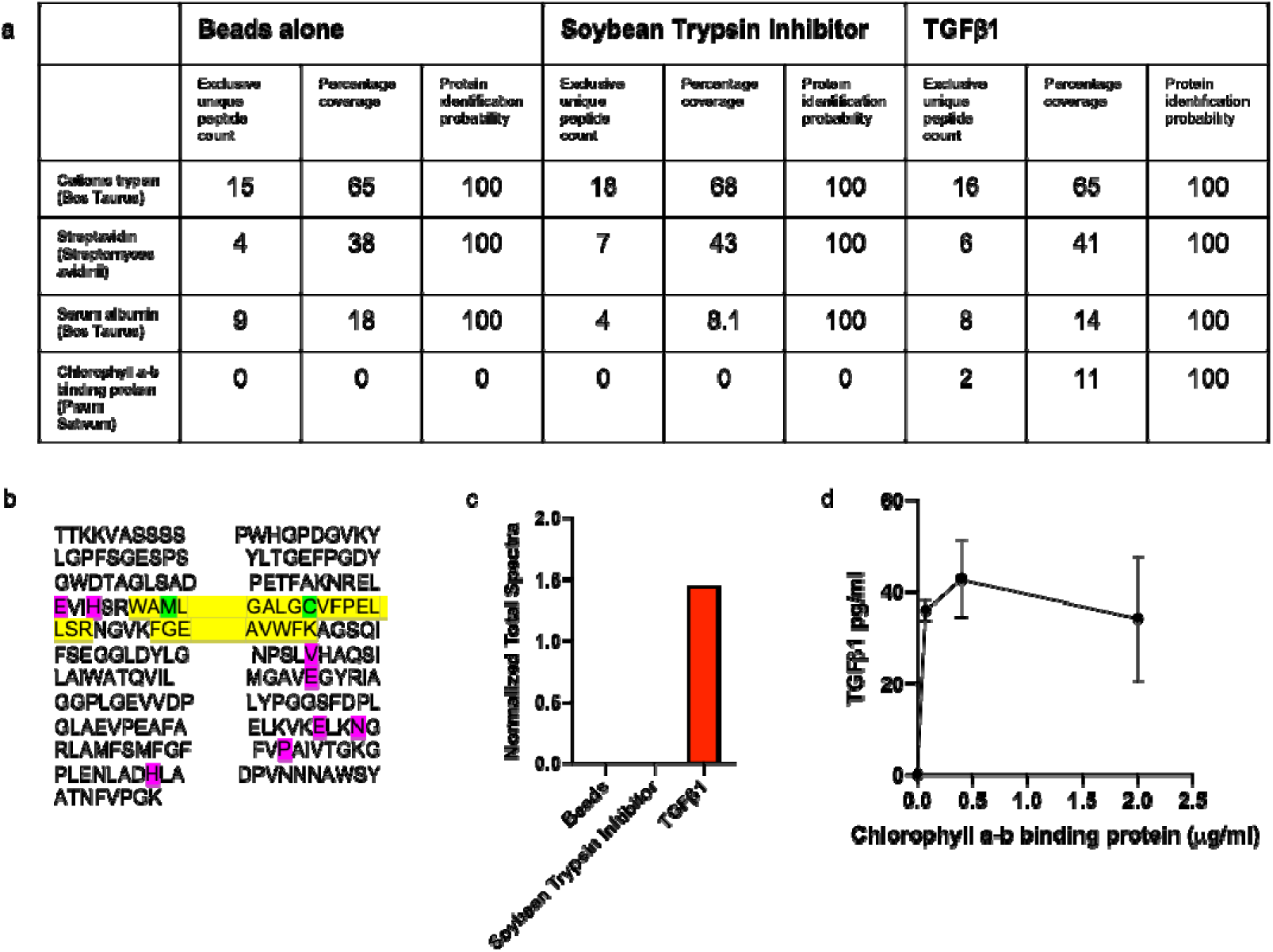
Chlorophyll a-b binding protein AB96 binds to active TGFβ1. (a) Mass spectrometry data for pulled out proteins when using indicated bait proteins when a threshold of min number of peptides =2, protein threshold =20% and peptide threshold =95%. (b) The chlorophyll a-b binding protein AB96 peptides observed by mass spectrometry are highlighted in yellow. Green represents amino acid modifications: M, oxidation and C, carbamidomethyl. Pink represents Magnesium binding sites. (c) Quantitation of chlorophyll a-b binding protein AB96 pulled from VA extract using indicated bait protein. (d) Concentration of active TGFβ1 eluted from different concentrations of chlorophyll a-b binding protein AB96 after incubation for 2h at RT.

This study reveals the first plant-derived cytokine binding protein. The central role that active TGFβ1 plays in many of the pathologies VA and other medicinal plants are ingested for, suggests that this protein, or a peptide derived from this protein, may help to explain some of the observed effects. It also opens the door for further discoveries where natural proteins/peptides alleviate pathology via the removal of immune evasion mechanisms, rather than innate stimulation or direct effects on the disease-causing agent.

One of our chief concerns with this work is the fact that monocots do not appear to be able to bind active TGFβ1 well, despite expressing Ca-bBP. We believe that this may be due the increased amount of vascular tissue per unit surface area compared to eudicots, with recommendations in the field that chlorophyll measurements between the two classes to be kept separate ^19^. Given our extract concentrations were based on total amounts we believe it would be reasonable to conclude our results were due to this much lower proportion of Ca-bBP in the starting material.

Ca-bBP AB96 is a component of the light harvesting complex of photosystem II, and as such it captures and delivers excitation energy. It binds at least 14 chlorophylls and carotenoids such as lutein and neoxanthin, as well as magnesium. Ca-bBP AB96 is membrane bound and difficult to isolate without detergents and sonication. It is possible that the endotoxin cleanup (detergent, vortexing and highspeed centrifugation) allowed for its presence in the extracts, indeed, with one of the peptides overlapping a transmembrane region (<u>WAMLGALGCVFPELLSR</u>; underlined region shows overlap) we believe that this must have occurred.

Although we have shown that the whole recombinant *E.coli* produced protein can bind TGFβ1, we do not yet know if the whole protein is required for this binding. The peptides picked up in the mass spectrometry are next to each other, and may be suggestive of a functional region, however further studies are required to confirm whether or not this is the case.

The question remains as to why Ca-bBP AB96 has the ability to bind active TGFβ1. We feel the answer may lie in relative charges; active TGFβ1 is more positively charged (isoelectric point; 7.73) and this charge is critical for its functionality in maintaining latency, whilst Ca-bBP AB96 is more negatively charged (isoelectric point; 4.88) enabling it to bind magnesium ions (along with chlorophyll, lutein and neoxanthin). This would make Ca-bBP AB96 a likely heparin-mimetic, acting as a scaffold to cause the oligomerization of TGFβ1 ^17^.

In terms of whether Ca-bBP AB96 has a real impact on malaria and other internal pathologies, this depends on the protein, or peptide’s, functional survival after acidification, neutralisation and digestion in the gastrointestinal tract, and entry into the circulation. Our mass spectrometry data show that the protein can be degraded by trypsin, however there is currently limited evidence of plant peptides entering and remaining in the circulation ^20^. For intestinal helminth infections, there are fewer delivery concerns as Ca-bBP AB96 peptides could act directly, or close to, the site of pathology without gaining entry into the circulation.

Regarding the reported efficacy of VA extract itself, the answer may lie in a combination of agents that act either directly on the pathogen (e.g. sesquiterpene lactones) or indirectly via immune polarization (e.g.β-glucans) or removal of immune evasion (Ca-bBP AB96). We now know enough about pathogenic mutation under selection pressure to understand that a single agent is unlikely to be efficacious for the length of time VA appears to have been used.

In conclusion, Ca-bBP AB96 has been shown to bind to and inactivate TGFβ1. The next step will be to identify whether this is a property of the whole protein, or merely a peptide fragment. In the case that this activity can be localized, then it may be possible to make a mimetic or a peptide which carries the same activity, and this would have potential for therapeutic intervention.

## Methods

### Preparation of leaf extracts

Defined species were sourced from UK and Ugandan suppliers. Fresh leaves were pulped with 50ml distilled water using a pestle and mortar. The extracts were filtered (45nm), before being boiled for 30mins and re-filtered (45nm). Samples were then subjected to three endotoxin clean-ups [until the LAL test proved negative], using Triton-X114 (Sigma) as described ^21^. The samples were then quantified either using a spectrophotometer or BCA assay (Pierce). Endotoxin tests (LAL; Lonza) were carried out in the presence of a β-glucan block (Lonza) [β-glucans present in plant cell walls cross react in the LAL assay] and samples were only used if levels were <0.1EU/mg.

### Generation of monocyte-derived dendritic cells

Monocytes were isolated from fresh blood packs (NHSBTS) using CD14 microbeads (Miltenyi Biotec) and the MACS system (Mitenyi Biotec). After treatment with 1500U/ml IL4 (BioTechne) and 400U/ml GMCSF (BioTechne), cells were cultured in AIM-V media (GIBCO) for 6 days. Cells were confirmed to be immature DCs using flow cytometry as described^22^. Extract was diluted in PBS to reach stock concentration for use on cells. Heparin sulphate (Sigma) was dissolved in PBS before use. Cell viability was assayed using Annexin-V/PI (BD biosciences) staining by flow cytometry.

### Protein binding assay

Leaf extract (1mg/ml in PBS), TGFβ1 antibody (5μg/ml in PBS; clone 1D11 BioTechne), parthenium (1mg/ml in chloroform; Sigma), artemisinin (1mg/ml in chloroform; Sigma), full length *Pisum sativum* L. (Leguminosae) chlorophyll a-b binding protein AB96 (at indicated concentrations; mybiosource TTKKVASSSSPWHGPDGVKYLGPFSGESPSYLTGEFPGDYGWDTAGLSADPETFAKNRELEVIHSRWAMLGALG CVFPELLSRNGVKFGEAVWFKAGSQIFSEGGLDYLGNPSLVHAQSILAIWATQVILMGAVEGYRIAGGPLGEVVDPLYPGGSFDPLGLAEVPEAFAELKVKELKNGRLAMFSMFGFFVPAIVTGKGPLENLADHLADPVNNNAWSYATNFVPGK) or vehicle (PBS or chloroform) were plated on a high binding 96 well flat-bottomed plate (Greiner) and left at RT overnight. The next day the plate was washed x3 in PBS + 0.05% Tween-20 (Sigma) before being blocked with 5% Tween-20 in PBS. Active TGFβ1 (BioTechne), IL-10 (BioTechne), IL-12p70 (BioTechne) or latent TGFβ1 (BioTechne) were added at indicated concentrations before being incubated for 2h at RT. The supernatant was removed and placed immediately in a relevant ELISA for assaying; this formed the ‘unbound fraction’. The plate was washed x3 in PBS + 0.05% Tween-20 before being treated with 50μl 1M HCl for 30mins at RT to elute any bound protein. This ‘bound fraction’ was neutralised using 50μl 1.2M NaOH/0.5M HEPES and immediately assayed in a relevant ELISA.

### ELISAs

TGFβ1, IL-12 and IL-10 ELISAs (all BioTechne) were performed as per the manufacturer’s instructions.

### TGFβ1 reporter assay

MINK cells (generously provided by Professor Rifkin) were cultured and assays performed as described^23^. Antibodies and VA extract were diluted in PBS prior to use. Cell viability was assayed using trypan blue after the assay period.

### Anion Exchange Chromatography

Samples were prepared as follows: 10ml 100ng/ml active TGFβ1 (BioTechne) in PBS. 10ml 50mg/ml VA extract in PBS. 5ml 100ng/ml active TGFβ1 in PBS plus 5ml 50mg/ml VA extract in PBS, mixed for 4h on a rotator at RT. Samples were loaded using a Superloop and run on an ACTApurifier 10 system using the equipment, reagents and programme described^24^. Fractions were assayed for TGFβ1 using an ELISA as described.

### Modified Immunoprecipitation

500μl 1μg/ml biotinylated active TGFβ1 in PBS (BioTechne), 500μl 1μg/ml biotinylated Soybean Trypsin Inhibitor (Biotechne) and PBS were each mixed with 500μl of 50mg/ml VA extract in PBS on a rotator for 4h at RT. 50μl magnetic streptavidin beads (Pierce) were washed in TBS + 0.1% Tween-20, added to each mixture, and mixing continued for a further 2h. Beads were pulled out using a magnetic stand and samples were washed x3 in TBS + 0.1% Tween-20. Samples were eluted from beads using SDS-PAGE reducing sample buffer (ThermoFisher) and run on SDS PAGE (ThermoFisher). Gels were stained and destained using a silver staining for mass spectrometry kit (Pierce). Whole lanes were cut out and analysed using mass spectrometry.

### Mass spectrometry

Gel sections were digested with bovine trypsin (Sigma) following reduction and alkylation of cysteine bonds with dithiothreitol and iodoacetamide (Sigma). Peptide extracts were subjected to chromatographic separation by C18 reversed-phase nano-trapping (Acclaim PepMap100 C18 Trap, 5 mm × 300 μm) and nano-analytical columns (EASY-Spray PepMap® C18, 2μm 100 Å, 75μm × 15cm) on an EASY NanoLC system (ThermoFisher) using a three-step linear gradient at a flowrate of 250nl/min over 60 minutes. The eluate was ionized by electrospray ionization using an Orbitrap Velos Pro (ThermoFisher) operating under Xcalibur v2.2. Precursor ions were selected according to their intensity using the collision-induced fragmentation employing a Top20 CID method. Raw spectral data was processed using Proteome Discoverer (v1.4) against the Uniprot ‘All Taxonomy’ database under the Mascot 2.2 algorithm (Matrix Science). Results were analysed using Scaffold Software (version 4.11.0; Proteome Software).

## Acknowledgements

Professor Joy Burchell (KCL) for allowing this research to be carried out in her laboratory and support. Professor Brian de Sousa (LSHTM/UCL) for advice and strategic input. Dr Sandrine Sellam (BioTechne) for advice and providing reagents. Professor Simon Croft (LSHTM) and Professor Adrian Hayday (KCL/CRICK) for support. Professor Daniel Rifkin (NYU) for the use of the TGFβ1 reporter cell line. Finally, Kato Sailus. K and Wasswa Drake. D for sharing their knowledge of this plant, inspiring this project and providing materials.

## Conflicts of interest

None

## Notes

### Competing Interest Statement

The authors have declared no competing interest.

